# Parameterization of Proximal Humerus Locking Plate Impingement with In Vitro, In Silico, and In Vivo Techniques

**DOI:** 10.1101/368258

**Authors:** Emily M. Bachner, Elaine C Schmidt, Matthew Chin, Surena Namdari, Josh R. Baxter, Michael W. Hast

## Abstract

**Background:** Locked plating of displaced proximal humerus fractures is common, but rates of subacromial impingement remain high. Computational predictions of implant impingement have yet to be sufficiently explored in proximal humerus fixation. The goal of this study was to utilize a multidisciplinary approach to elucidate the relationships between common surgical parameters, anatomical variability, and the likelihood of plate impingement.

**Methods:** The experiment was completed in three phases. First, a controlled *in vitro* experiment was conducted to simulate impingement. Second, a dynamic *in silico* musculoskeletal model was developed to simulate changes to implant geometry, surgical techniques, and acromial anatomy, where a collision detection algorithm was used to simulate contact between the plate and acromion. Finally, *in vivo* shoulder kinematics were recorded for nine activities of daily living and motions that created a high likelihood of impingement were identified.

**Results:** Impingement was measured at 73.3±14.5° abduction in the cadaveric model and 92.0°±34.0° with computational simulations. Impingement events were limited to ranges of motion between 10–40° of cross-body adduction. Activities of daily living, such as combing one’s hair, lifting and object overhead, and reaching behind one’s head are likely to cause impingement.

**Discussion and Conclusion:** This multidisciplinary experiment quantified key preoperative factors to assist with implantation decisions. Results demonstrated that proximal implant placement, superior translation of the humeral center of rotation, increases in plate thickness, and increases in acromial tilt all increase the likelihood of impingement. Careful preoperative planning that includes these factors could help guide operative decision making and improve clinical outcomes.

Level of Evidence: V

## Introduction

Proximal humeral fractures have become the third most common fracture type in patients over 65 years of age^4,20,27,30^ and are expected to increase three-fold over the next 30 years.^1^ The development of locking plate technology for proximal humeral fracture fixation has become increasingly used and widely accepted, particularly for patients with osteoporotic bone.^2^ Unfortunately, reported complication rates are high, ranging from 20% to 49% in some studies.^8,11,26^

Impingement between the acromion and the proximal portion of a locking plate is believed to be a potential source of pain that may also limit the patient’s range of motion. Preventing post-operative subacromial impingement is especially difficult due to wide variability in shoulder joint morphology within humans.^21^ One study has reported that 42.4% of patients opted to remove hardware due to impingement related issues.^19^ Previous studies have suggested that proximal positioning of the plate may lead to impingement^5,28^, and there are many reasonable causes for such a scenario. Proximal plate positioning may be due to slight errors in surgical technique, purposeful positioning in order to optimize fixation, or as a result of small humerus anatomy paired with a single-sized implant. To date, no studies have attempted to biomechanically quantify “how high is too high?” or provided quantifiable guidelines with respect to implant design, shoulder anatomy, and desired range of motion when considering subacromial impingement.

The objective of this study was to systematically quantify locking plate-subacromial impingement with an array of variables associated with proximal humerus fixation and anatomical variation. The study used a combination of *in vitro, in silico, and in vivo* models to characterize post-operative function of the joint. Accurate estimations of surgical impingement, caused by changes in surgical and anatomical parameters, will help provide clarity on this complicated issue.

## Materials and Methods

This experiment was performed in three phases (Figure 1). First, a controlled dynamic cadaveric model was created to record impingement events during simple abduction motions. Second, a validated musculoskeletal model of the upper extremity was used to identify the onset of subacromial impingement. In this computational setting, a series of controlled abduction motions were simulated while variables of plate placement, acromial geometry, and humerus center of rotation (COR) were systematically changed. Third, kinematics associated with nine activities of daily living (ADLs) in healthy patients were recorded with 3-D motion capture. Shoulder joint angles were compared to simulation outputs and high risk ADLs were identified.

**Figure 1:**
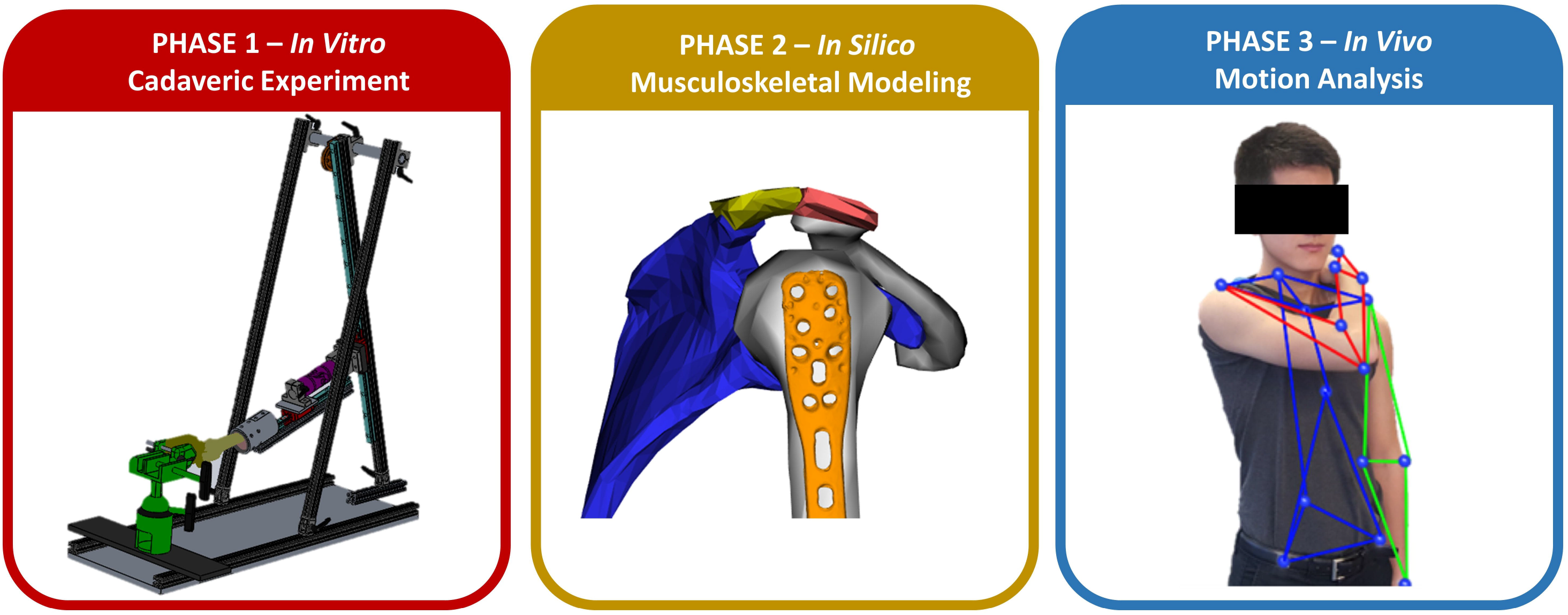
A workflow diagram outlining the methods used to perform the study. Impingement events were recorded during simple abduction motions in Phase 1. Computational simulations of prescribed motions with systematically changing variables were made in Phase 2. Phase 3 utilized motion capture techniques to compare shoulder joint angles during activities of daily living to those output in Phase 2.

### Phase 1 – In Vitro Experiment

Four cadaveric upper extremities (2M, 2F, mean age 66.75 years) were used in this study. Shoulder joints were isolated and the humerus was amputated at the midshaft. All unnecessary soft tissue was removed from each specimen and the distal humerus was potted in polycarbonate tubing with polymethyl methacrylate (Lang Dental, Wheeling, IL). Locking plates (Philos, DePuy Synthes, West Chester, PA) were securely fixed on the humeri per manufacturer guidelines with two bicortical screws (3.5 mm diameter, 35 mm length).^10^ A fracture was not simulated and no screws were inserted into the humeral head because this study focused solely on impingement and not fixation strength. Specimens were imaged with fluoroscopy (SIREMOBIL Compact (L) C-Arm, Siemens, Washington, DC) in anterior/posterior and medial/lateral outlet views. Post-hoc measurements were made (ImageJ, National Institutes of Health, Bethesda, MD and Matlab, Mathworks, Natick, MA) for humeral head radii, acromial tilt (the angle between the antero-inferior edge of the acromion, postero-inferior edge of the acromion, and the inferior tip of the coracoid process), and acromial slope (the supplement of the angle between the postero-inferior edge, the inferior aspect, and antero-inferior edge the acromion) (Supplemental Figure 1).^21^

Several preparatory steps were required to track joint kinematics and detect subacromial impingement in real time. For the purpose of motion capture, retroreflective marker clusters were rigidly attached to the plate and scapula. Thin-film pressure sensors (iScan 6900, Tekscan, Inc., South Boston, MA) were covered with a patch of cellophane tape that extended beyond the boundaries of the sensor. Beaded Kirschner wires were inserted through the tape and into the proximal portion of the humerus, such that the sensor consistently covered the proximal aspect of the plate during motions (Supplemental Figure 2).

Simulations of abduction were created with a custom-built jig (Figure 2). Scapulae were held stable with a scapula clamp (Pacific Research Laboratories, Vashone, WA) and a vise. The potted distal humeri were fixed to a sled that allowed free translation on a low-friction linear bearing. This assembly was attached to a second sled via a universal joint. The second sled was allowed to translate on a linear bearing that was angled 60° relative to horizontal. A rope and pulley connected the sled to the rotational actuator of a universal test frame (ElectroForce 5500, TA Instruments, New Castle, DE). Initial shoulder angles were controlled by rotating the scapula in the vise and re-clamping, and dynamic simulations of 20° of abduction were performed in 5 seconds. Adjustments were made to the initial scapular position within the vise until an impingement event was created.

**Figure 2:**
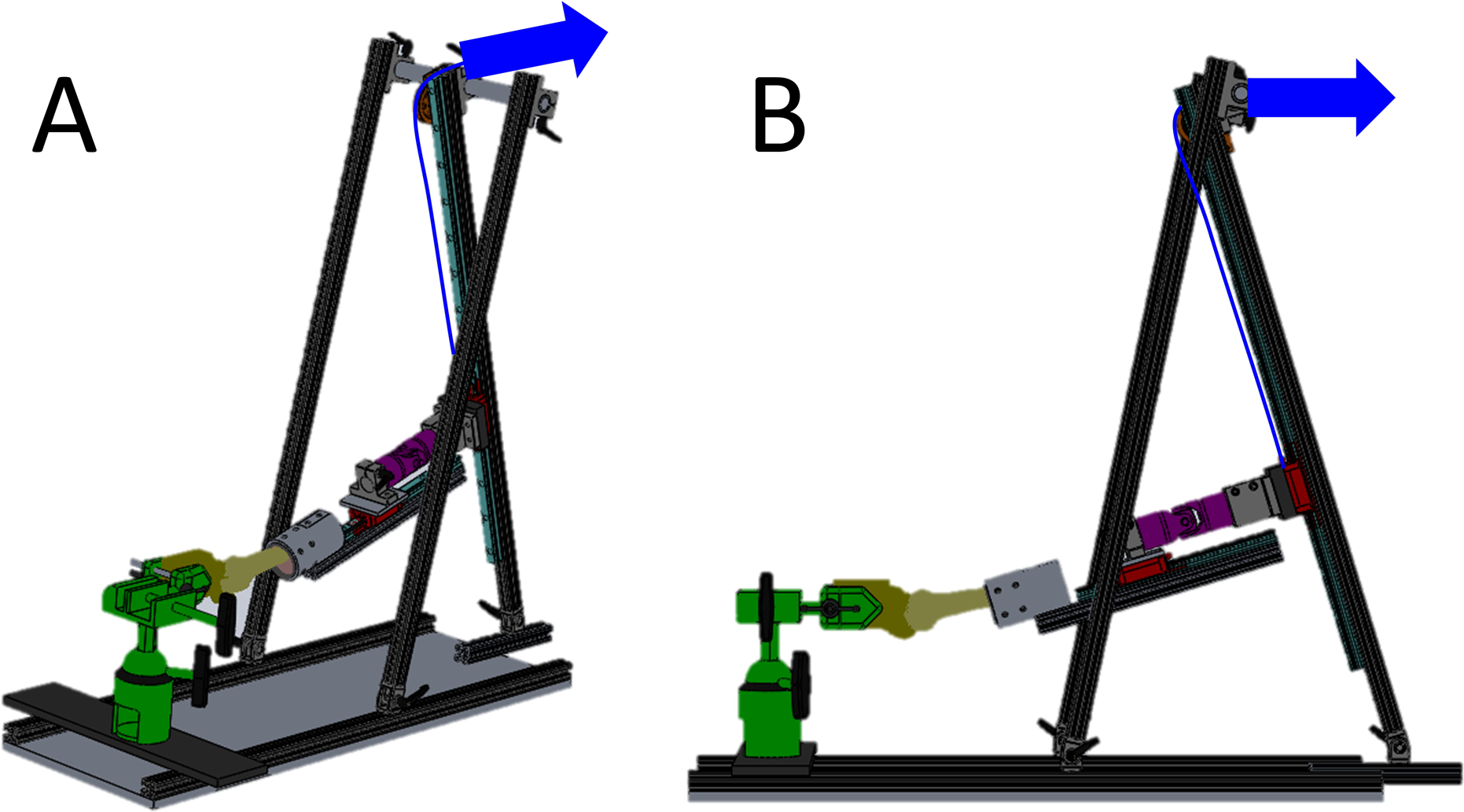
Computer-aided drawings of the custom jig built for the experiment in isometric (A) and side (B) views. The scapulae were held in place with a scapula clamp attached to a vise (green). Two sleds (red) were attached to one another with a universal joint (magenta), and the sleds were allowed to freely translate on linear tracks (teal). Displacement of the rope (blue) created an abduction motion, controlled by a rotational actuator of a universal test frame (not shown).

Scapular and humeral motions were tracked in real time using a 6-camera motion capture system (Optitrack, NaturalPoint, Inc., Corvallis, OR) calibrated to 0.2 mm accuracy. Five anatomic points of the scapula were identified with an instrumented wand during static trials so that the post-hoc computational model could be properly scaled (Supplemental Figure 3). Motions of the marker clusters were recorded as the shoulder was abducted until impingement occurred.

To determine scapulohumeral joint angles, simplified representations of each cadaveric humerus and scapula were created in a simple 2 body, 6 degree-of-freedom, OpenSim model.^9^ The on-board geometry files of the humerus and scapula were scaled to represent the cadaveric specimens (See Appendix A for details). A 3-D rendering of the locking plate was created by performing an optical 3-D scan (Afinia Einscan, Afinia, Chanhassen, MN). The virtual plate was placed on the humerus in the model per manufacturer guidelines and held in place with a weld joint. Anatomic coordinate frames were assigned to the humerus and scapula, and marker trajectories were tracked with the inverse kinematics algorithm. The relative 3-D motions between the virtual humerus and scapula were characterized as a function of time. The timed data from the pressure sensor measurements were synchronized with the marker trajectory data, and initiation of impingement was identified by distinct increases in compressive forces.

### Phase 2 – In Silico Experiment

A validated computational model of the upper extremity of a 50^th^ percentile male was adapted for this phase of the experiment.^25^ Use of this model permitted the simulation of motions that are not easily recreated in a cadaveric setting and provided the ability to make systematic and controlled changes to other variables of interest. Shoulder kinematics were defined using a validated spherical coordinate system.^16^ This mechanical convention ensures the smooth execution of complex shoulder motions, but it utilizes nomenclature that is substantially different than clinical standards. For the purpose of clarity, results will be described in the clinically relevant terms of cross-body adduction (humeral motion in the transverse plane) coupled with abduction (humeral elevation in the coronal plane).

Surgical and anatomical variables were introduced into the model to identify possible impingement events (Figure 3). Position of the plate on the humerus was varied from a neutral location using two parameters: 1) proximal-distal displacements (−10 to +10mm in 5mm increments) and 2) plate thickness (0 mm, 2.5mm, and 5.0mm). Acromial tilt and slope were changed between 20–35° in 5° increments. Finally, the humeral head COR ranged from neutral to +5.0mm proximal in 2.5mm increments. This step was performed because the model utilizes a joint that does not translate during motions. Moving the COR proximally, relative to the scapula, represents translations that may occur during a motion.^7^

**Figure 3:**
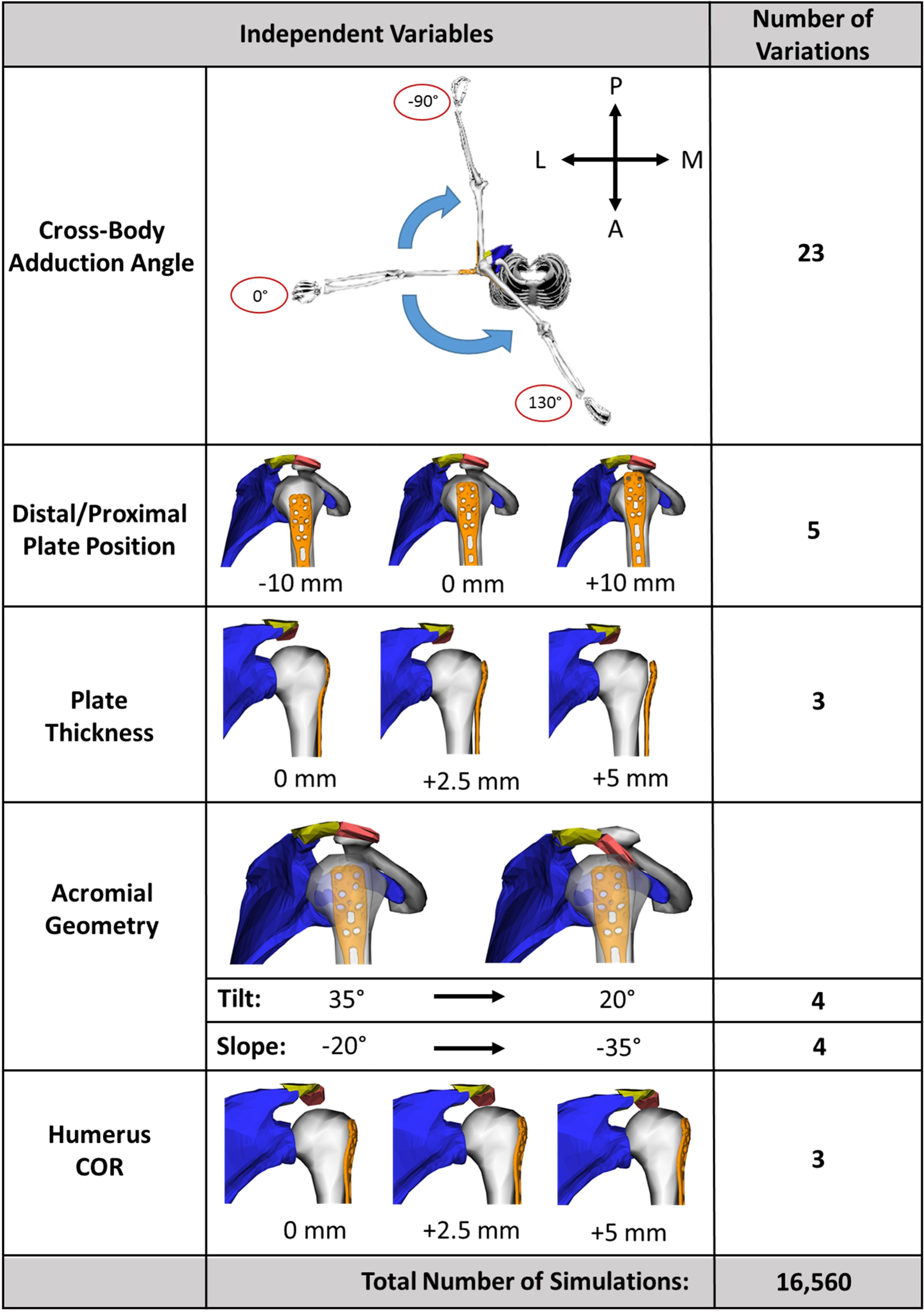
A diagram outlining the combinations of simulations that were performed with the in silico model.

Twenty-three unique simulations were executed to cover the entire range of motion of the shoulder joint (Figure 3). For each simulation, the initial cross-body adduction angle was systematically adjusted from –90° to 130° in 10° increments. With this pose established, the arm was lowered to 0° abduction and raised to180°. The on-board OpenSim elastic foundation contact algorithm was used to detect impingement between the plate and the acromion. Shoulder joint angles were recorded at the moment these collisions occurred. Results were pooled, and the parameters that were present during impingement were identified.

### Phase 3 – In Vivo Experiment

Eight healthy young subjects with no history of shoulder injuries or pain (4 M, 4 F, mean age 21.5 years) performed nine commonly performed ADLs^23^ after providing written informed consent in this IRB approved study. Upper extremity kinematics were measured using a 12-camera motion capture system (Raptor Series, Motion Analysis Corp, Santa Rosa, CA). Reflective markers (9.5mm, B&L Engineering, Santa Ana, CA) were adhered bilaterally using skin-safe tape covering the 7^th^ cervical vertebra, sternum, acromion, elbow epicondyles, and ulnar and radial styloid processes. Subjects stood upright with their arms straight and shoulders at 90° abduction and external rotation to scale the subject-specific musculoskeletal models. Marker labeling was visually confirmed, gaps were filled using cubic-spline interpolation, and marker trajectories were filtered. Shoulder kinematics were calculated using the same musculoskeletal model used in Phase 2. Analysis of the root mean squared, total squared, and maximum error indicated that this model created was a good representation of the experimental kinematics. Using a boot-strapping technique, 95% confidence intervals for elevation angle, abduction, and internal rotation were calculated for each ADL. Results were compared to computational predictions of impingement based on joint angle.

## Results

### Phase 1 – Cadaveric Experiment

The cadaveric experiment measures were made in terms of scapulohumeral angulation. Simulated impingement occurred at a mean cross-body adduction angle of 22.1 ± 10.1°, abduction angle of 73.3 ± 14.5°, and external rotation of 26.7 ± 13.4°. Mean humeral head radii were 23.1±2.4 mm. Acromial geometry identified using fluoroscopic imaging showed mean acromial tilt of 26.2°±3.3° and acromial slope of 27.6 °±5.7°.

### Phase 2 – Computational Model

Computational output measures were made in terms of thoracohumeral angles. Impingement only occurred when cross-body adduction angles were set between 10° to 50°, with a mean of 31.4°±9.6°. (Figure 4A). More than one-in-ten simulations of this range of motion, (368 out of 3,600 simulations) exhibited impingement, which occurred between 56° and 178° of abduction, with an average of 92.0°±34.0° (Figure 4B). Although it is possible to have impingement when the plate is placed distally (18.4%), 73% of impingement events occurred when the plate was moved proximally beyond the neutral location (Table 1). Similarly, increases in plate thickness led to increases in impingement events. Decreases in acromial tilt led to higher rates of impingement, with 84% of impingement events occurring when tilt was set to either 20° or 25°. Changes in acromial slope had no impact on the likelihood of impingement. Finally, proximal shifts of the humeral head COR also led to increases in impingement. For the sake of simplicity, these results have been presented on a variable-by-variable basis, but, complex relationships exist within this data set. For more information regarding these relationships, the reader is referred to the Appendix B in the supplementary material.

**Table 1.**
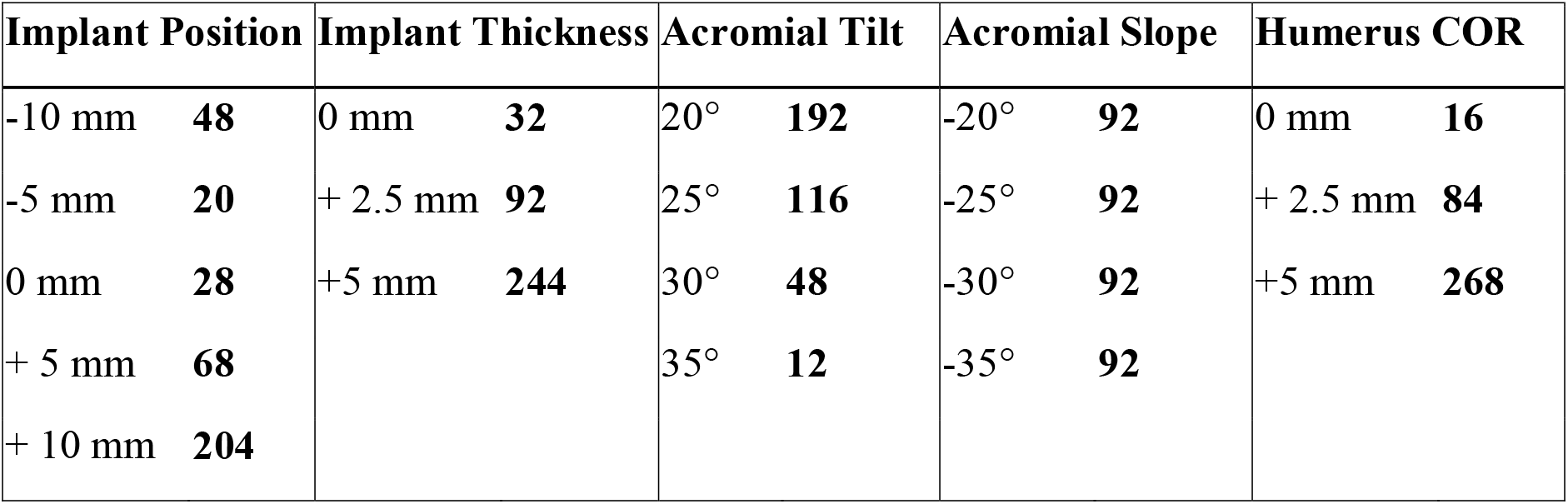
Breakdown of computational model results showing number of impingement events per modified parameter.

| **Implant Position** |  | **Implant Thickness** |  | **Acromial Tilt** |  | **Acromial Slope** |  | **Humerus COR** |  |
| --- | --- | --- | --- | --- | --- | --- | --- | --- | --- |
| -10 mm | **48** | 0 mm | **32** | 20° | **192** | -20° | **92** | 0 mm | **16** |
| -5 mm | **20** | + 2.5 mm | **92** | 25° | **116** | -25° | **92** | + 2.5 mm | **84** |
| 0 mm | **28** | +5 mm | **244** | 30° | **48** | -30° | **92** | +5 mm | **268** |
| + 5 mm | **68** |  |  | 35° | **12** | -35° | **92** |  |  |
| + 10 mm | **204** |  |  |  |  |  |  |  |  |

**Figure 4:**
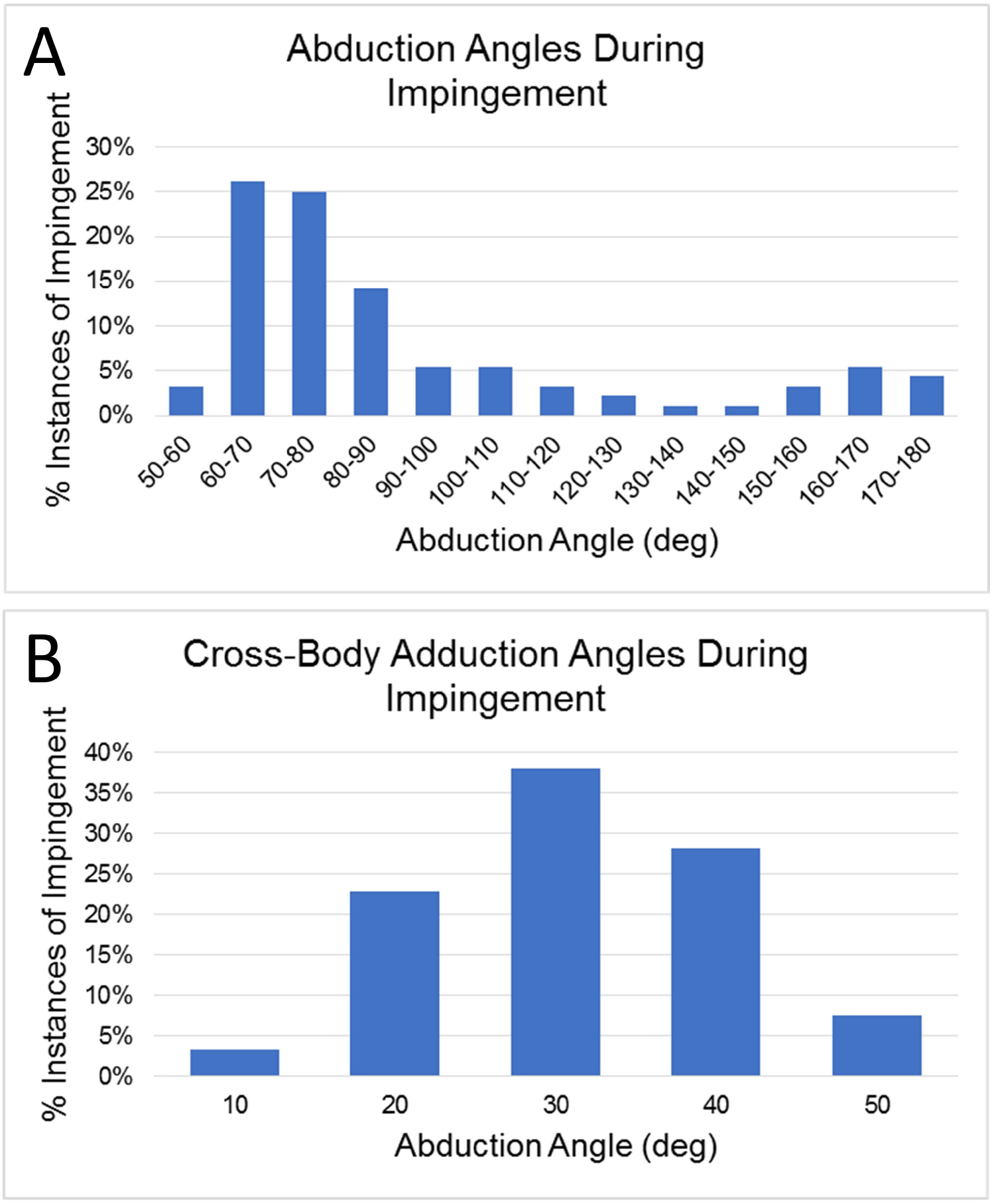
Histograms showing the distribution of the impingement simulations with respect to cross-body adduction angle (A) and abduction angle (B).

### Phase 3 – In Vivo Motion Comparison

Of the 9 activities of daily living that were recorded, only 3 motions produced shoulder kinematics in which impingement occurred within the model. Comparisons between the *in silico* and *in vivo* data revealed that reaching behind the head, lifting a light object overhead, and combing hair are activities with a high likelihood of impingement (Figure 5). Other motions created joint angle combinations that did not create impingement in the simulations (Supplemental Figure 4).

**Figure 5:**
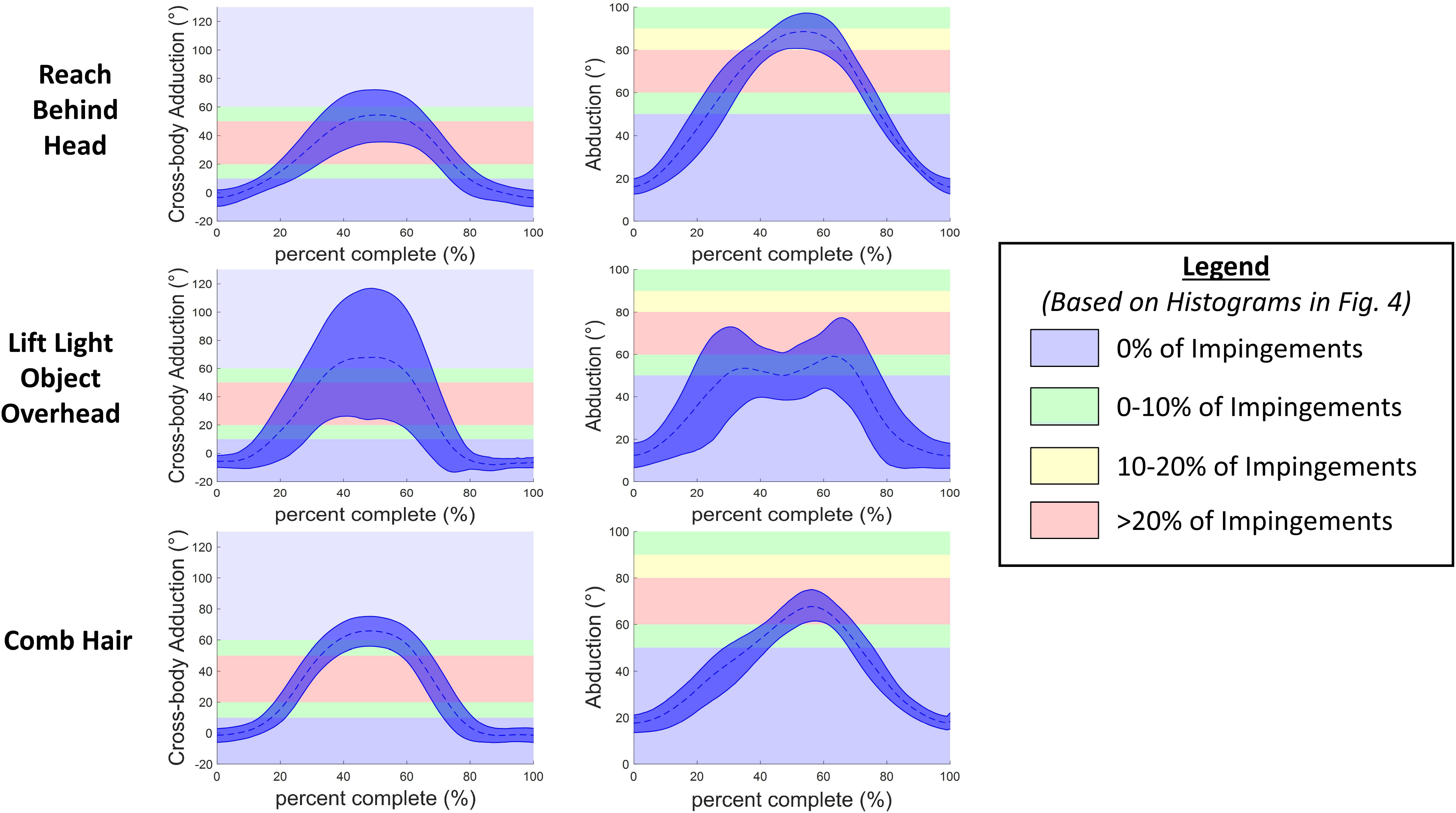
Plots of cross-body adduction (left column) and abduction (right column) during activities of daily motion. Mean values (blue dashed) and +/-one standard deviation (blue cloud) are shown. Plots are overlaid on backgrounds that are colored to represent the number of impingements reported in the histograms in Figure 4.

## Discussion

This study improves the biomechanical understanding of locking plate-subacromial impingement and the findings compare favorably to previous *in vivo* and *in vitro* studies. A previous cadaveric experiment measured an average glenohumeral impingement angle of 74°±15°^18^, which is very similar to the 73.3 ± 14.5° found in Phase 1 of this experiment. Results from the computational model matched well with static MRI studies that investigated changes in subacromial space.^12–14^ Dynamic evaluations of subacromial impingement using open MRI techniques found impingement at 93.5° of thoracohumeral abduction in asymptomatic patients.^29^ This *in vivo* outcome matches well with the 92.0°±34.0° simulated in the current study. The computational analysis in Phase 2 utilized a generalized 50^th^ percentile male model. The use of this size provided a reasonable approximation of human anatomy, based on the small number of cadaveric specimens used in Phase 1. Specifically, the radius of the humeral head (23.5 mm) and the ranges of acromial slopes and tilts (20°-35°) fell within the range of the measurements taken in the cadaveric specimens (23.1±2.4 mm, 27.6±5.7°, 26.2±3.3° respectively).

Differences between the cadaveric model and the computational model can be attributed to the use of scapulohumeral angles in the cadaveric model (due to a lack of a thorax), while thoracohumeral angles are used in the *in silico* and *in vivo* models. It is tempting to believe that the higher abduction angles observed in the computational model indicate a later onset of impingement. Interestingly, the opposite is true. When the scapular rhythm is accounted for, the mean scapulohumeral joint angles (73.3° abduction, 22.1° cross-body adduction) occur when the arm is positioned at approximately 118° of thoracohumeral abduction and 40° cross-body adduction (Figure 6). This over approximation of thoracohumeral impingement joint kinematics in the cadaveric model is likely due to a lax capsule, which may have caused the humeral head to move posteriorly relative to the glenoid.

**Figure 6:**
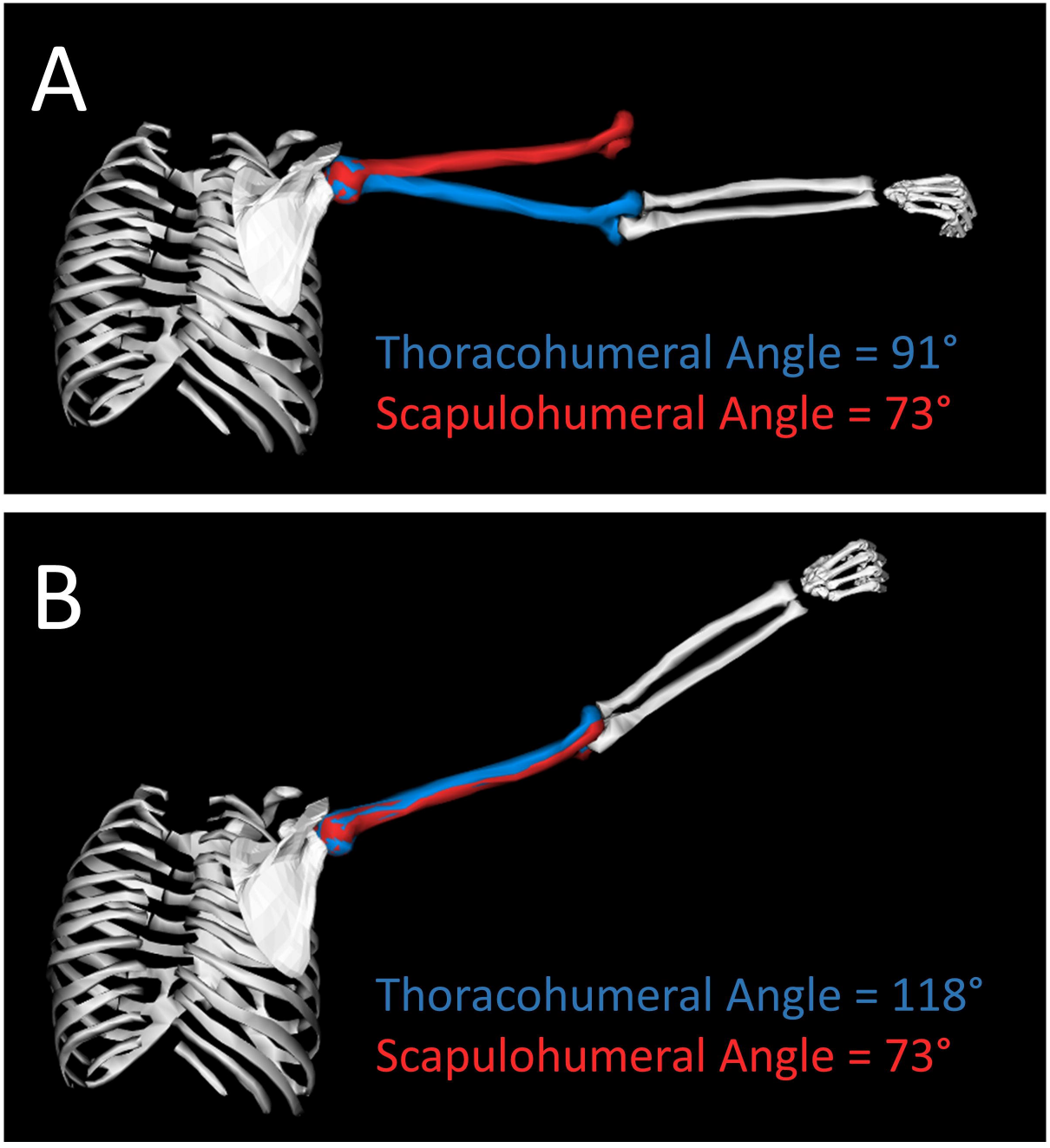
Images from a musculoskeletal model that superimposes results from cadaveric (red) and computational (blue) models (A). When scapular rhythm is accounted for, the arm must be in approximately 118 degrees of thoracohumeral abduction to achieve 73 degrees of glenohumeral abduction.

The cadaveric jig developed in this experiment is unique. Existing abduction simulators often consist of a large, custom-made, semi-circular ring to guide abduction motions.^3,17^ This existing design provides the ability to simulate muscle-driven motions over large ranges of motion; however, it also makes the rig both expensive and cumbersome. The rig developed in the current study provides a smooth and continuous shoulder motion. The range of motion can be changed by reorienting the scapula at the beginning of a test. The created motions are passive in nature, and the weight of the sled must be considered. This rig was developed with linear, off-the-shelf, components for less than $1000 and can fold down to size for easy storage.

The musculoskeletal model developed in this study is also noteworthy. A variety of previous shoulder simulators have been developed to estimate muscle loads during motions and to investigate implant stresses and strains.^6,15,24,31^ To our knowledge, this model represents the first attempt to create a model that is tailored specifically to change surgical and anatomic variables related to locking plate impingement. Changes to 4 of the 5 anatomic and surgical variables resulted in differences in subacromial impingement timing. Interestingly, changes to acromial slope did not have an effect on the model. As mentioned previously, the acromial slope is determined by calculating the supplement of the angle between the postero-inferior edge, the inferior aspect, and antero-inferior edge the acromion. In order to make an adjustable acromial slope within the model, a “hinge joint” was placed at the inferior aspect of the acromion. Post-hoc analysis of simulation outputs show that collisions between the plate and the acromion always occurred in the area posterior to the location of this “joint.”

This experiment has several limitations. The cadaveric simulations represent only passive motions, and were not driven by coordinated muscle activations. Changes to the model’s basic boney geometry may alter simulation results. Since the subacromial space is relatively small, even a subtle change in any one of these variables could potentially result in changes to the onset of impingement. This is important to keep in mind, given that subacromial space dimensions are highly variable due to factors including sex, muscle activity, acromion morphology, posture, and age.^12,22^ Aside from translating the humeral head COR, translations in the glenohumeral joint were constrained, which may not fully characterize the human condition. Simulation output suggests that impingement does not appear to be sensitive to variation in acromial slope. However, a previous study compared patients with impingement syndrome to controls and found no significant difference in acromial slope between the two groups.^21^ Therefore, this finding may have clinical relevance, and this topic requires further investigation. Movement biomechanics may differ between patients following a surgical repair of a proximal humerus fracture and the healthy-young adults with no history of upper extremity injury that participated in this study. However, the framework of this study supports the concepts that patient anatomy, surgical placement, and hardware parameters all can affect subacromial impingement risks.

## Conclusion

Open reduction internal fixation of proximal humerus fractures have relatively high complication rates, some of which can be attributed to subacromial impingement. Results from this experiment suggest that patient anatomy in conjunction with implant characteristics could help guide operative decision making. This study successfully implemented a multidisciplinary workflow that utilized *in vitro* biomechanical experimentation, *in silico* musculoskeletal modeling, and *in vivo* 3-D motion capture to quantify subacromial impingement. Results from this experiment quantified key preoperative factors to assist with implantation decisions. It also confirms the importance of accounting for scapular rhythm and glenohumeral stability when simulating impingement. The data from the current experiment provides valuable information to clinicians and rehabilitative specialists to better predict patient outcomes and guide rehabilitation. Future studies within patient populations may help predict the likelihood of subacromial impingement and identify post-surgical activity guidelines that may further reduce complication rates and improve overall outcomes.

**Author Contribution Summary**
**EMB:** Contributed to research design, acquisition, analysis, and interpretation of data. Drafted and revised the paper. Read and approved the final submitted manuscript.
**MC:** Contributed to research design, acquisition, analysis, and interpretation of data. Drafted and revised the paper. Read and approved the final submitted manuscript.
**ECS:** Contributed to research design, acquisition, analysis, and interpretation of data. Drafted and revised the paper. Read and approved the final submitted manuscript.
**SN:** Contributed to research design and interpretation of data. Provided revisions of the paper. Read and approved the final submitted manuscript.
**JRB:** Contributed to research design, acquisition, analysis and interpretation of data. Drafted and revised the paper. Read and approved the final submitted manuscript.
**MWH:** Contributed to research design, acquisition, analysis and interpretation of data. Drafted and revised the paper. Read and approved the final submitted manuscript.

## Acknowledgments

The authors would also like to thank Anthony Cresap for his help with cadaveric testing and Todd Hullfish and Annelise Slater for their help collecting motion capture data.

